# An auditory advantage of *Rdl*-resistant mosquitoes may promote its persistence in urban environments

**DOI:** 10.64898/2026.05.20.726456

**Authors:** C Sangbakembi-Ngounou, J Bagi, E Doran, Y-G Gonetomy, J Mbiko-Tanza, S Tytheridge, C Quinn, D Ayala, T Nolan, M Andrés

## Abstract

Despite dieldrin being discontinued, resistance mutations in the GABA-gated chloride channel RDL (*Resistance to dieldrin*) continue at high frequencies in natural malaria mosquitoes, suggesting a selective advantage. We showed that RDL modulates the mosquito auditory sensitivity. Because acoustic recognition is essential for mate acquisition, we hypothesized that resistance mutations confer a pleiotropic effect on mating success, with potential consequences for sexual selection. We provide laboratory evidence that resistance mutations enhance auditory and mating behaviours. We then conduct field collections in the city of Bangui (Central African Republic) and rural areas, revealing an association between *Rdl* resistance genotypes and urban settings, and within the city, with the noisiest locations. We also show decreased mating success of susceptible females with increasing noise levels, suggesting detrimental effects. Together, our findings support that *Rdl* resistance mutations enhance auditory function and mating success. We propose that this auditory advantage may contribute, together with other selective pressures such as cross-selection by other insecticides, to the persistence of these alleles in nature and may facilitate urban colonization by malaria vectors.

## Results and discussion

Mosquitoes recognize their mating partners through hearing their flight tones. Their ears are unique across insects because of their anatomical complexity and their high functional plasticity, which is mediated by an extensive efferent neuromodulatory network ^1–3^ (Fig. 1A). We recently described a role of the GABA_A_ receptor *Rdl* (*Resistance to dieldrin*) in modulating the auditory sensitivity of the malaria mosquito *Anopheles gambiae*. We showed that blocking RDL with picrotoxin, a well-characterised antagonist, leads to increased spontaneous activity of the antennal nerve and enhanced auditory responses ^1,4^, suggesting a role for this gene in mating behaviour. Similar effects have been described in *Culex quinquefasciatus*^1,2^, suggesting high conservation of GABAergic auditory inhibition across mosquito species. *Rdl* was first described as the locus where target-site mutations to the insecticide dieldrin were located^5,6^, primarily an alanine to serine (A296S) or glycine (A296G) in the channel’s second transmembrane domain ^7,8^. Dieldrin was widely used for malaria control in Africa from the 1950s to the 1970s as an indoor residual spraying (IRS) insecticide, when it was discontinued due to widespread resistance, high toxicity and environmental persistence^7–9^. Despite the continent-wide discontinuation of dieldrin over four decades ago, *Rdl* resistance alleles persist at substantial, and in some cases increasing, frequencies across the *An. gambiae* complex ^9,10^. Classical evolutionary theory predicts that resistance alleles should decline after insecticide withdrawal due to associated fitness costs^10^, but this has not been the case for *Rdl* resistance mutations upon dieldrin cessation without a clear explanation ^11^. Because hearing is essential for mosquito mating, we investigated if a RDL auditory role could affect mating-related fitness and thereby contribute to the persistence of resistance mutations in natural mosquito populations.

**Figure 1:**
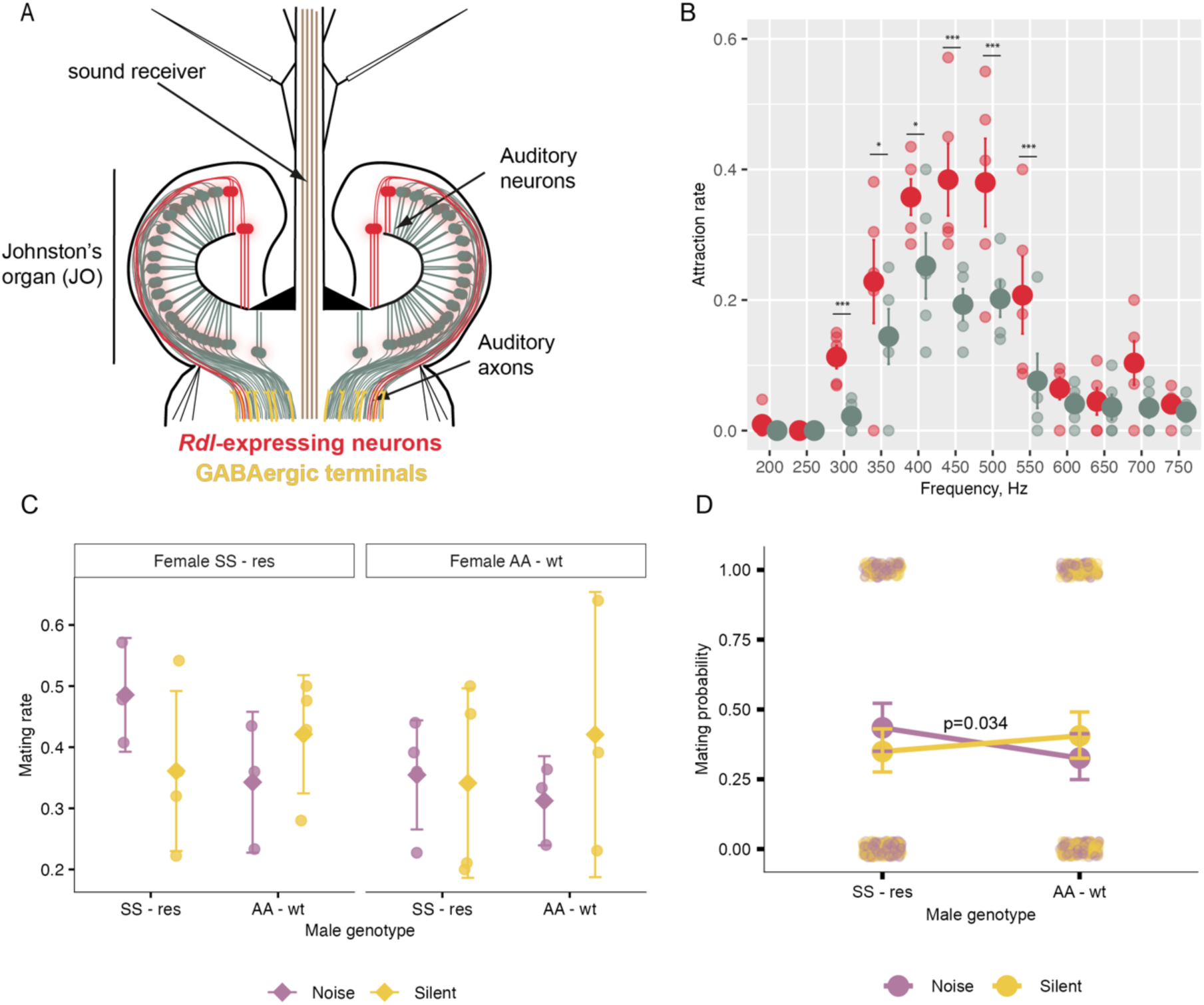
Effects on auditory and mating behaviours of A296S *Rdl* resistance alleles. **A**) Schematic of a male mosquito *An. gambiae* ear, including the sound receiver (flagellum), and the auditory organ or Johnston’s organ (JO) located at its base. The JO harbours *Rdl*-expressing auditory neurons^4^, and receives GABAergic innervation from efferent neurons in the auditory nerve^4^. **B**) Male mosquitoes were exposed to pure tones of different frequencies (200-750 Hz, at 50Hz intervals) and attraction rates to the sound source (phonotactic behaviour) were assessed. Homozygous A296S males (SS - res) were more attracted to the sound source compared to control WT males (AA - wt) (logistic regression, p-value<2e-16, graph shows post-hoc comparisons of genotype within each frequency). Each large point represents a replicate of 20 males and error bars represent ±1 standard error (SE). **C**) Mating rate across experimental treatments. Points represent replicatelevel proportions of inseminated females (number inseminated / total per replicate). Larger symbols indicate group means, and error bars show 95% confidence intervals calculated across replicates. Data are shown for each combination of male genotype, female mosquito group, and treatment. **D)** Predicted mating probability as a function of male genotype under low-noise (70 dB) and high-noise (93 dB) acoustic conditions. Jittered points show individual mating outcomes. Solid lines and filled points represent model-predicted probabilities from a generalized linear mixed model (GLMM) with a binomial error distribution and logit link, including treatment × male genotype interaction, treatment × female genotype interaction, and replicate as fixed effects, with date included as a random effect. Error bars represent 95% confidence intervals around model predictions on the probability scale. Predictions were marginalised over female mosquito genotype. Lines connect fitted values across male genotypes within each acoustic treatment. A significant interaction between male genotype and environmental noise treatment was detected (binomial GLMM, p = 0.034), indicating that the effect of noise on female insemination depends on male genotype.

We first assessed auditory behaviours in the laboratory of a dieldrin resistant Tiefora strain of *Anopheles coluzzi*. The strain was not fixed for the A296S *Rdl* mutation so following molecular characterization we generated a homozygous resistant (homozygous for the A296S mutation and therefore referred to as SS resistant, kindly provided by Linta Grigoraki, Liverpool School Tropical Medicine) and a homozygous susceptible line (referred to as WT, see Methods). These two lines were compared in laboratory behavioural assays.

Male mosquitoes are attracted to and fly towards sound sources that mimic female flight tones (phonotactic behaviour), reflecting their courtship behaviour ^12–15^. We analysed phonotactic responses of male mosquitoes in the lab by assessing attraction rates to a speaker broadcasting different pure tones to mimic female wing beat frequencies (sine waves, 200-750 Hz covering the mosquito auditory range, at 50 Hz intervals ^16^) (Fig. 1B). Homozygous SS resistant males consistently showed higher response rates than WT males to frequencies in the range of 300-550Hz, which are the most relevant biologically corresponding to female flight tones^16^ (logistic regression, p-value<2e-16). This result suggests that resistant mosquitoes have an enhanced auditory sensitivity.

We then analysed the mating behaviour of resistant and WT mosquitoes in the lab. We conducted experiments under two noise conditions, namely a “high-noise” incubator (93 dB) and a “low-noise” one (70dB; the incubators produced this background noise levels due to their machinery and fan systems, it was therefore technically impossible to test lower environmental noise levels) and performed different crosses of resistant and WT individuals (Fig. 1C-D). We modelled the results using a generalized linear mixed model (GLMM, binomial), including treatment, male mosquito number, and female mosquito number with interaction terms (treatment × male genotype, treatment × female genotype), replicate as a fixed effect, and date as a random intercept. A significant interaction between treatment and male genotype was detected (GLMM, β = 0.687 ± 0.329, z = 2.08, p = 0.037), indicating that the effects of environmental noise on female insemination rates differed depending on the male genotype. No significant interaction was found between treatment and female genotype. Previous studies have shown that sound masking can interfere with mosquito hearing ^17,18^. The increased sensitivity of mutant individuals shown in the phonotactic assays seemed to compensate for the detrimental effects of noise in mating partner detection.

Taken together our laboratory results including 1) increased phonotactic responses of resistant male mosquitoes, and 2) an interaction between noise levels and male genotype in female insemination rates, suggest that *Rdl* insecticide resistance mutations may confer an auditory advantage that increases mating success in noisy conditions. Given the persistence of *Rdl* resistance mutations in natural malaria mosquito populations, we hypothesized that the auditory advantage of mutant individuals could contribute to sexual selection. We therefore planned to collect evidence from the field in Central African Republic (CAR), where different species of the *An. gambiae* complex co-exist, and insecticide resistance profiles, including *Rdl* resistance mutations have been reported^19^. Bangui, the capital, and the surrounding rural areas, where anthropogenic noise levels are expected to vary, offer a natural setting to test our hypothesis.

### Field collections reveal urban enrichment of *Rdl* resistance alleles

We first assessed the ecology of *Rdl* resistance mutations in CAR. We collected *An. gambiae* complex mosquitoes from 7 urban sites in the city of Bangui (Central African Republic) and 4 surrounding rural sites (Fig. 2A) using both larval and adult early morning sampling (Supp.Table 1). Overall, *An. coluzzii* was most abundant in urban sites, whereas *An. gambiae s.s.* dominated rural areas (Fig. 2B). The odds of *An. gambiae s.s* presence in the city was 0.37 compared to *An. coluzzii* (Fisher’s exact test, OR = 0.37, p = 1.79e-06) consistent with known ecological preferences^20^. Hybrid individuals were also detected (6.7%) (Table 1, Fig. 2B). We examined the presence of *Rdl* resistance mutations (A296S/G, referred to as AS or AG) and observed 12.7% mutant genotypes across all samples collected, all in heterozygosity. Different studies have reported fitness costs of homozygous *Rdl^R^* mosquitoes, but not heterozygous individuals ^21,22^. *Rdl* resistance mutations in heterozygosity confer intermediate resistance to dieldrin exposure ^23,24^.

**Figure 2:**
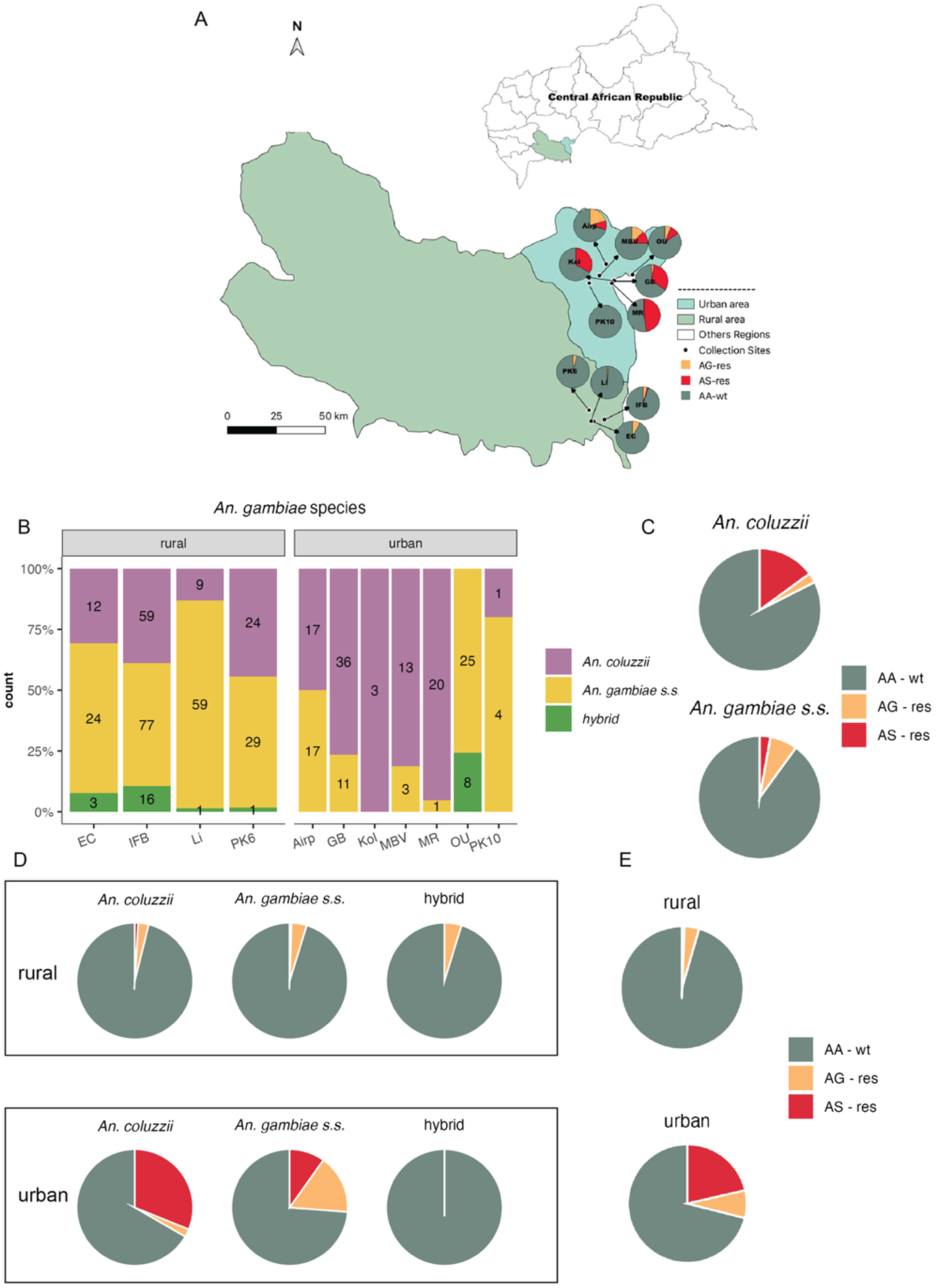
*Rdl* genotype distribution in the city of Bangui and rural areas. **A)** Mosquitoes were collected from larval sites and early morning indoor resting sites in seven urban locations in the city of Bangui and four rural villages. **B)** Composition of *An. gambiae* complex mosquito species collected in each site (including both males and females), showing *An. coluzzii, An. gambiae* and hybrids. Overall *An. coluzzii* was most abundant in cities, and *An. gambiae* in rural sites (Fisher’s exact test, p = 1.79e-06, OR = 0.37, 95% CI: 0.24 – 0.57). **C)** *Rdl* genotype distribution (susceptible wild-type as AA; A296G heterozygote as AS-res; A296S heterozygote as AG - res) by species. *An. coluzzii* showed higher overall resistance compared to *An. gambiae* (Fisher’s exact test, OR = 1.91, p = 0.024, 95% CI: 1.06 – 3.48). **D)** Resistant genotypes by species and location show urban enrichment of resistant genotypes. **E)** Data summarizing *Rdl* genotype distribution by location show that resistant genotypes were more frequent in urban areas, in particular AS heterozygotes were massively enriched in urban compared to rural sites (Cochran-Mantel-Haenszel test stratified by species, OR= 35.9, p= 1.7×10⁻¹¹).

**Table 1:** Distribution of resistant *Rdl* genotypes by zone and species (both larval and early morning collections).

| Site area | Species | N | AA – wt<br>n | AG - res<br>n (%) | AS - res<br>n (%) | Total resistant<br>n (%) |
| --- | --- | --- | --- | --- | --- | --- |
| Urban | *An. coluzzii* | 90 | 60 | 2 (2.2%) | 28 (31.1%) | 30 (33.3%) |
|  | *An. gambiae* | 61 | 45 | 10 (16.4%) | 6 (9.8%) | 16 (26.2%) |
|  | *Hybrid* | 8 |  | 0 (0%) | 0 (0%) | 0 (0%) |
|  | Total Urban | 159 | 113 | 12 (7.5%) | 34 (21.4%) | 46 (28.9%) |
| Rural | *An. coluzzii* | 104 | 100 | 3 (2.9%) | 1 (1.0%) | 4 (3.8%) |
|  | *An. gambiae* | 189 | 180 | 8 (4.2%) | 1 (0.5%) | 9 (4.8%) |
|  | *Hybrid* | 21 |  | 1 (4.8%) | 0 (0%) | 1 (4.8%) |
|  | Total Rural | 314 | 300 | 12 (3.8%) | 2 (0.6%) | 14 (4.5%) |

Overall, *An. coluzzii* showed higher rates of resistance compared to *An. gambiae* (Fig. 2C, Table 1, 17.5% vs 10.0% Fisher’s exact test, OR = 1.91, p = 0.024). Moreover, *An. coluzzii* presented higher frequencies of AS compared to AG (15% vs 2.6%, Fisher’s exact test, OR = 7.22, 95% CI: 2.47 – 28.83 p= 1.549e-05), while the opposite was observed for *An. gambiae* (2.8% vs 7.2%, 18/250; Fisher’s exact test, OR = 0.38, p = 0.04), as previously described^25,26^. We did not observe differences across sexes (Fisher’s exact test, OR = 0.73, p = 0.25).

When analysing the spatial distribution of mutant genotypes across locations, we observed that resistance mutations were enriched in urban (28.9%) compared to rural (4.5%) sites (Fig. 2D, Fisher’s exact test, OR = 8.68, p = 2.795e-13). Moreover, analysis of individual genotypes revealed dramatically different patterns in their distribution (Table 1-2, Fig. 2D). The AS genotype was massively enriched in urban environments: 21.4% frequency in urban compared to only 0.6% (2/314) in rural specimens after controlling for species (Fig, 2E, Table 1-2, Cochran-Mantel-Haenszel test stratified by species, OR= 35.9, p= 1.703e-11). AG also showed an urban enrichment (7.5% in urban vs 3.8% in rural mosquitoes), but the association was weaker than for AS (Cochran-Mantel-Haenszel test, OR = 2.53, p = 0.033). Our results show an enrichment of *Rdl* resistance genotypes in urban locations, suggesting a selection of *Rdl* resistance alleles in the urban context.

### Resistant genotypes are enriched in the noisiest locations within the city of Bangui

Anthropogenic noise is one of the main urban pollutants affecting wildlife ^27^. To uncover potential association between anthropogenic noise and *Rdl* resistance mutations, we measured environmental noise levels around our collection sites. As expected, noise measurements (Fig. 3A, Supp. Table 2) revealed significantly higher levels in urban (mean = 61.6 ± 8.5 dBA, n = 7) compared to rural sites (mean = 49.5 ± 4.1 dBA, n = 4) (Welch two-sample t-test, t = -2.78, df = 9.59, p = 0.02). This confirms an association between urban settings and anthropogenic noise^28^, although noise patterns were complex in Bangui (Fig. 3A).

**Fig. 3:**
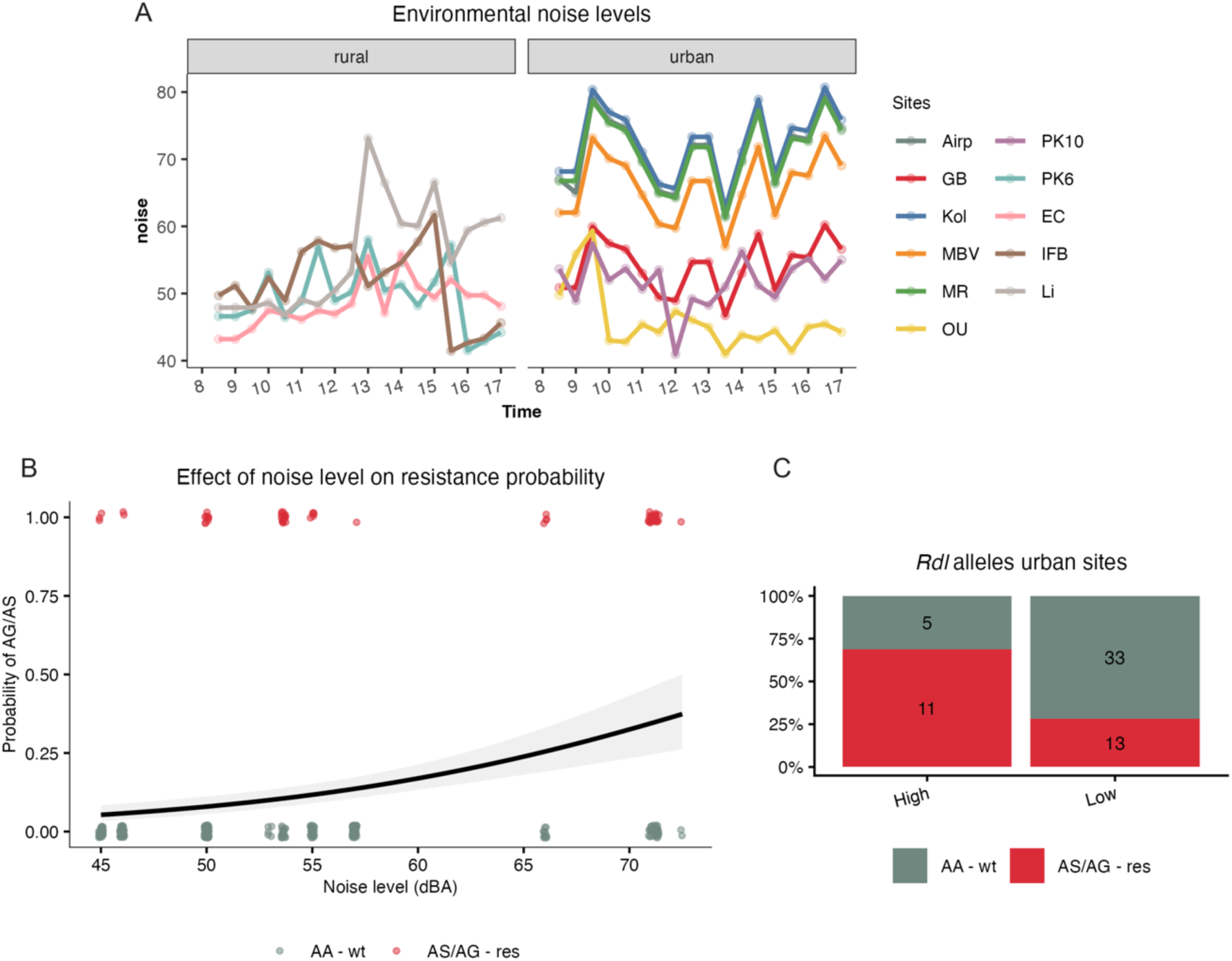
*Rdl* resistance mutations are associated with higher environmental noise levels. **A**) Noise levels in the different study locations along the day, including urban and rural areas. Overall, noise levels were higher in the city (Welch two-sample t-test, t = -2.78, df = 9.59, p = 0.02). **B**) Effect of environmental noise level on *Rdl* resistance probability. Logistic relationship between noise level and resistance probability. Points represent individual observations (coloured by genotype, AA wild-type vs. AS/AG resistant), with a fitted binomial GLM shown by the solid line and 95% confidence interval. Each 1 dBA increase in noise level was associated with a 9.0% increase in the odds of resistance (OR = 1.09, p = 3.17e-08, 95% CI: 1.06–1.12), suggesting that noise may act as a selective pressure. **C**) Analysis of genotype distribution across the two noisiest and the two quietest urban locations shows resistant genotypes are enriched in the noisiest urban locations (Fisher’s exact test: p = 0.007, OR = 5.41, 95% CI: 1.37–30.68).

**Table 2:** Assessment of the association between *Rdl* resistance mutations (data pooled for both species) and urban location compared to rural settings by estimating the odds ratios and the statistical significance, using Cochran-Mantel-Haenszel test stratified by species for individual mutations and Fisher’s exact test for both mutations together.

| Phenotype | *Rdl* genotype | urban | rural | OR |
| --- | --- | --- | --- | --- |
| WT | AA -wt | 113 | 300 |  |
| Resistant | AG - res | 12 | 12 | AG-urban vs AG-rural<br>OR = 2.53, p = 0.033 |
|  | AS -res | 34 | 2 | AS-urban vs AS-rural<br>OR= 35.9, p= 1.703e-11 |
|  | AG/AS - res | 46 | 14 | AG/AS-urban vs AG/AS -rural<br>OR = 8.68, p = 2.795e-13 |

We first tested the association between mutant genotypes and environmental noise levels, independently of whether locations were rural or urban (Fig. 3B). Logistic regression analysis revealed a significant positive association between ambient noise levels and the presence of resistant mutations. For every 1 dBA increase in noise, the odds of an individual mosquito being resistant increased by approximately 9% (OR = 1.09, p = 3.17e-08). This provides evidence linking anthropogenic noise to the distribution of *Rdl* resistance mutations.

As noise levels were variable across urban sites (Fig. 3A), we attempted to study the association between anthropogenic noise and *Rdl* genotypes within the urban ecosystem; however, the resulting model was underpowered and did not allow reliable inference. Instead, we compared the odds of mutant genotype presence (pooling data from both species) between the two noisiest urban sites (“Kol”, mean noise level = 72.5 dB; “MR” = 71 dB) and the two quietest urban sites (“OU” = 46 dB; “PK10” = 51.8 dB). Mutant genotypes were significantly enriched in the noisiest locations (Fig. 3C; Kol: 33%, MR: 48%, OU: 15%, PK10: 0%; Fisher’s exact test: p = 0.007, OR = 5.41, 95% CI: 1.37–30.68), further supporting the core hypothesis within the urban landscape itself.

### Field Mating Success

We further investigated differences in the insemination status of susceptible and resistant females collected in early morning sampling (Fig. 4). Interestingly, while in rural conditions 75% of susceptible AA females were mated, in urban environments only 45% of susceptible females carried sperm, depicting a positive association between rural environment and susceptible female mating success (Fig. 4A-B, Fisher’s Exact test, OR=3.65, p=0.004). This result may suggest a difficulty of susceptible females to mate in the urban context. We further explored this correlation by studying the relationship between susceptible female mating success and environmental noise independently of the location (Fig. 4B). Logistic regression modelling confirmed a decrease of approximately 13% in the probability of insemination for each 1 dBA increase in noise levels. By contrast, when considering *Rdl* resistant alleles, all AS/AG mutant females captured in the city were mated, although their numbers were low (4 AS - res and 2 AG - res females). In rural settings, only two AG mated females were captured and 1/2 was mated. Low sample size prevented meaningful statistical comparisons, but these data suggest a pattern of higher insemination rates of resistant females in the city that highlights the need of further field studies with increased sample sizes.

**Figure 4:**
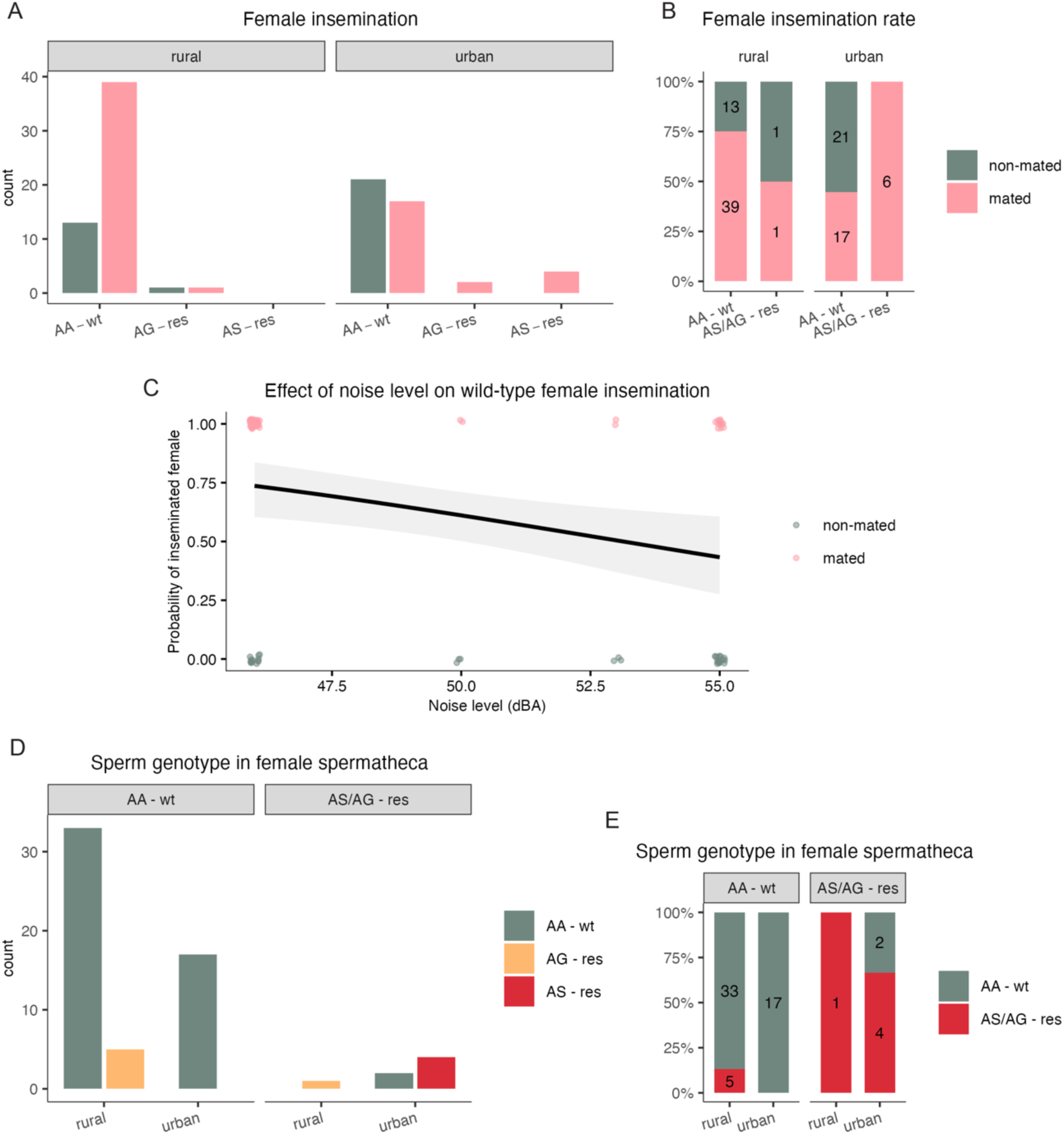
*Rdl* genotypes and female insemination in field mosquitoes. **A**) *Rdl* genotype of female mosquitoes collected in rural and urban environments. Susceptible wild-type females were more likely to be inseminated in rural settings (Fisher’s Exact test, p=0.004, OR=3.65). *Rdl* mutant females collected in urban areas were all mated, while only 1/2 (50%) was mated in the rural area. **B**) Summary of female insemination rates pulling data for both *Rdl* mutant genotypes shows that susceptible females were more likely to be mated in rural environments. By contrast, heterozygous resistant females were more commonly mated in the city, although statistical analysis was not possible due to low sample size. **C**) Effect of environmental noise level on mating success of susceptible wild-type females. Points represent individual observations (coloured by insemination status), with a fitted binomial GLM shown by the solid line and 95% confidence interval. Each 1 dBA increase in noise level corresponded to an approximate 13% reduction in the probability of insemination (OR = 0.87, p = 0.07, 95% CI: 0.78– 0.96), suggesting that noise may reduce successful mating of wild-type females. **D**) Analysis of *Rdl* genotypes of sperm collected from spermatheca dissected from mated females. Counts are shown grouped by female genotype. **E**) Sperm genotype proportions between female genotypes in rural and urban areas. Although sperm genotypes do not differ across AA susceptible females in rural and urban locations (Fisher’s Exact test, p=0.31), AS/AG resistant females appear to mate assortatively with mutant males, although low sample numbers prevent statistical comparisons.

A causal association between the reduction in susceptible female mating rates and environment noise is difficult to prove in field studies. It is important to note that current knowledge implies a predominant role of males in acoustically detecting mosquito females ^29^, while the function that hearing plays in *An. gambiae* female courtship is unknown ^30^. In this regard, the fact that our field data show that noise affects susceptible female mating is intriguing. To disentangle this, we also analysed the sperm genotype of collected females to infer the *Rdl* genotype of the male that successfully mated with them. Grouping by female genotype reveals no statistical difference in genotypic composition of sperm across susceptible AA females collected in rural and urban sites (Fig. 4 D-E, Fisher’s Exact test, p=0.31, the estimated odds ratio was infinite because no resistant sperm genotype was observed in urban females). By contrast, the frequency of mutant genotypes was higher across AS/AG resistant females in both rural and urban locations (Fig. 4C), however, low sample sizes prevented meaningful statistical comparisons. These data suggest in any case assortative mating mechanisms, which have been shown before^31^. This would explain why even if *Rdl* resistance mutations where not affecting female hearing, assortative mating would favour mutant females in the city as *Rdl* genotypes are more common in this setting at the mosquito population level. It is also plausible that *Rdl* resistance mutations also enhance female hearing, increasing their mating success in noisy environments. In any case, validating these hypotheses requires further investigation.

## Conclusion

The persistence of *Rdl* resistance mutations in natural mosquito populations decades after dieldrin cessation is a puzzling phenomenon that has intrigued the scientific community for years ^11^. This presence has been associated with cross-resistance mechanisms with other insecticides such as the pesticide fipronil ^32^, which is mainly used in agriculture^33^ and cattle protection ^34^. Interestingly, some studies from 1960 reported high incidence of dieldrin resistance in natural mosquito populations in Nigeria where dieldrin was never used before (and where other insecticides were used in a very limited scale)^35,36^, suggesting other type of selective advantage apart from survival to insecticide use. Our study supports that dieldrin resistant mosquitoes present an auditory advantage mediated through a direct role of RDL in mosquito hearing.

We previously showed that *Rdl* expressed by auditory neurons in the Johnston’s organ (JO) modulates the auditory sensitivity of *An. gambiae* through inhibiting the mosquito ear’s spontaneous activity and auditory responses to sound stimulation ^1,4^. Previous electrophysiological studies of *Xenopus laevis* oocytes expressing wild-type and mutant *Rdl* showed that resistance mutations affected GABA responses, suggesting an alteration of RDL sensitivity to GABA in mosquitoes carrying *Rdl* resistance mutations ^37^. In our laboratory studies, we show that homozygous A296S mosquitoes are more responsive to sound compared to susceptible individuals. Moreover, we also show an interaction between male genotype and noise treatment on mating success (Fig. 1). These data support the hypothesis of an auditory advantage.

Our field collections in Central African Republic, indicate an ecological association between *Rdl* resistance mutations and urban environments, and within the city, with the noisiest locations. Moreover, field collections of adult females suggest a deleterious effect of anthropogenic noise on susceptible female mating success. Together, our data suggest a mechanism of sexual selection where *Rdl* resistance mutations are kept in natural malaria mosquito populations through an advantage to acoustically detect mating partners. The selection of *Rdl* resistance mutations would strengthen urban adaptation of malaria mosquitoes by enhancing resilience to anthropogenic noise. This would exemplify a case of urban adaptation through exaptation ^38^, where a trait that evolved under historical insecticide selection is co-opted for a new function in a different ecological context^39^, facilitating urban colonization. Interestingly, we show that the distribution of AS and AG resistance mutations differs across settings, being A296S enriched in the city compared to A296G. Likewise, we and others^25,26^ have shown that AS are more frequent in *An. coluzzii* compared to *An. gambiae*. In this regard, it should be noted that different compensatory mutations are associated to resistance mutations in codon 296 in natural mosquito populations ^11^. These compensatory mutations have been shown to alter responses of RDL to GABA *in vitro* ^37^. Understanding effects of resistance haplotypes in RDL function and its behavioural implications could shed light on the mechanisms underlying the ecological differences observed.

Previous studies have shown higher frequencies of insecticide resistance alleles in cities ^39–41^, linked to pesticide and insecticide use for vector control, agriculture and farming ^42–44^. Although dieldrin is no longer used, *Rdl* resistance mutations have been shown to provide cross-resistance to the pesticide fipronil^32^ and have been related to the persistence of *Rdl* resistance mutations in natural mosquito populations. However, other forms of uncharacterized selective advantage have been proposed as the effects of cross-resistance mechanisms are not fully clear ^11^. We show that sexual mechanisms driven by an auditory advantage of *Rdl*-resistant individuals that increases its mating success could contribute to the maintenance of the high mutant allele frequencies observed in nature. Although our field study has clear limitations related to the complexity of inferring mechanisms from ecological associations or the difficulty of isolating noise effects from other potential urban confounders, such as light and chemical pollution, we provide evidence from behavioural data collected in the laboratory to support the causal relationship between noise and increase *Rdl*-resistant genotypes beyond ecological association.

In summary, we show that *Rdl* resistance mutations confer an auditory advantage in *An. gambiae* and provide ecological evidence to support that these auditory effects contribute to sexual selection to drive the persistence of mutant alleles in natural mosquito populations and adaptation of insecticide resistance mosquitoes to urban environments. This implies that insecticide selection can have far-reaching, unpredictable consequences, enhancing vector resilience to human ecosystems ^45^. This underscores the need for an “evolutionary mindset” in vector control^46^, anticipating how interventions might shape behavioural and ecological traits ^47^. As most insecticides are neurotoxic with broad effects on mosquito biology, monitoring not only resistance allele frequencies but also associated behavioural phenotypes will be crucial for predicting and mitigating the unintended evolutionary outcomes of interventions, ensuring the long-term sustainability of malaria control.

## 2. Methods

### 2.1. Mosquito Strains and Rearing for Behavioral Assays

An *An. gambiae* Tiefora line carrying A296S *Rdl* resistance mutations was used in the laboratory experiments. The line was not fixed for the mutations and therefore it was screened to isolate heterozygotes and wild-types using the LNA-rdl-296 assay (see below) to start the establishment of a A296S homozygous (A296S-Hom) and susceptible A296A (Tiefora) colonies for the experimental work. Mosquitoes were reared under standard insectary conditions (28 ± 1°C, 80 ± 10% RH, 12:12 light: dark cycle with one hour light transition at dawn and dusk) and provided 10% sucrose ad libitum. Virgin males and females were isolated at the pupal stage. Experiments were conducted with 3-to-5-day old mosquitoes.

### 2.2. Molecular Analyses

#### DNA extraction and species identification

Genomic DNA was extracted from individual mosquito specimens using Nexttec™ 1-Step DNA Isolation Plates (Nexttec Biotechnologie GmbH, Germany) following the manufacturer’s protocol for tissue samples. This method provides high-purity DNA suitable for downstream PCR applications. Species identification within the *Anopheles gambiae* complex was performed using a standard multiplex PCR assay distinguishing *An. gambiae* s.s., *An. coluzzii*, and *An. arabiensis* ^48^.

#### Rdl genotyping (LNA-rdl-296 assay)

We employed a quantitative PCR (qPCR) assay based on Locked Nucleic Acid (LNA) hydrolysis probes for high-resolution allelic discrimination at the *Rdl* codon 296 locus. This assay simultaneously detects the wild-type alanine (Ala) allele and the two major resistance alleles-serine (A296S, predominant in *An. coluzzii* and *An. arabiensis*) and glycine (A296G, predominant in *An. gambiae* s.s.) in a single reaction.

A 118-bp fragment surrounding the target codon was amplified using primers Rdl-296-1F (5’-TCGTGGGTATCATTTTGGCTA-3’) and Rdl-296-1R (5’-TTTTCGGTAAGGCAGCATTC-3’). Allele-specific detection was achieved with three LNA probes, each labeled with a distinct fluorophore:

- Rdl-296-Ala: HEX-labeled, specific for the wild-type allele (sequence: CA+CG+TGT+T+G+CATT-AGG).
- Rdl-296-Gly: 6-FAM-labeled, specific for the A296G resistance allele (sequence: CA+CG+TGT+T+G+GA+TTA).
- Rdl-296-Ser: Cy5-labeled, specific for the A296S resistance allele (sequence: AGCA+CG+TGT+T+T+CA+TTAG).

qPCR reactions were performed in 10 µL volumes containing 1× PrimeTime Gene Expression Master Mix (IDT), 0.2 µM of each primer, 0.1 µM of each LNA probe, and 1 µL of DNA template. Amplification was carried out on a QuantStudio or Aria MX real-time PCR system equipped with filters for HEX, FAM, and Cy5, using the following thermal profile: initial denaturation at 95°C for 3 min; 40 cycles of 95°C for 5 sec and 64°C for 30 sec.

Fluorescence data (ΔRn values) for each dye were exported. Due to platform software limitations for tri-allelic calling, genotypes were determined using a custom Microsoft Excel template by calculating the fluorescence signal above the automatically set threshold for each channel. A positive allele call was recorded when the ΔRn difference exceeded zero. Genotypes were assigned as homozygous (Ala/Ala, Ser/Ser, or Gly/Gly) or heterozygous (Ala/Ser or Ala/Gly) based on the combination of positive signals. Only samples with clear amplification curves and unambiguous allele calls were included in the final analysis.

### 2.3. Laboratory Mating Assays Under Acoustic Conditions

Mating assays were performed in custom cages (15 x 15 x 15 cm) placed inside one of two identical environmental incubators. One chamber served as the “low noise” condition, with a background noise level of 70 ± 2 dB SPL at the cage center. This was the baseline environmental level of the incubator, caused by the fan and other background components. Due to this equipment limitations, it was not possible for us to test lower environmental noise levels. The other was the “high noise” condition, where continuous white noise (0.1–20 kHz) was broadcast via a speaker to maintain a constant 93 ± 2 dB SPL at the cage center, simulating intense urban noise ^49^. The incubator had three outlined areas for the cages to sit and the cages were randomly assigned for each experiment to one of the three locations.

For each assay, 25 virgin males and 25 virgin females (3 to 5 day old) of specified genotypes were released into a cage at the onset of scotophase, and were exposed for the first time to the different environmental noise conditions. Three crosses were tested: i) Tiefora males x Tiefora females, ii) A296S-Hom males x Tiefora females, iii) Tiefora males x A296S-Hom females, iv) A296S-Hom males x A296S-Hom females. Mosquitoes were allowed to mate for 48h (instead of a single swarming period) to allow them to acclimatise to noise levels, females were removed, cold-anesthetized, and their spermatheca dissected in PBS and examined under 400x magnification for sperm presence. Each cross-condition combination was replicated three times. The examiner was blind to the genotype of mosquitoes being dissected.

### 2.4. Phonotaxis (Acoustic Attraction) Assays

Male attraction to sound was tested in a 15 x 15 x 15 cm cage located inside an environmental incubator. Twenty male mosquitoes were placed inside the cage immediately after eclosion and entrained to a 12:12 light:dark cycle with one hour light transition at dawn and dusk inside the incubator (ZT0-ZT1 and ZT12-ZT13, respectively; ZTX is the formalized notation of an entrained circadian cycle’s phase). At day 3, a speaker placed at the cage wall was used to broadcast pure tone sounds (200-750 Hz, at 50 Hz intervals, SPL are provided in Supp. Table 4, spanning the mosquito hearing range) of 1-minute duration at ZT13, when lights were completely off. Tone frequencies were randomised. The experimenter, who was blind to the mosquito genotype (Tiefora or A296S homozygoues), counted the number of mosquitoes landing on the speaker.

### 2.5. Study Area and Mosquito Collections

Fieldwork was conducted in urban locations in Bangui, capital of the Central African Republic (CAR) (4.3947° N, 18.5582° E), and rural locations across villages between June 2023 and March 2024. The area is hyperendemic for malaria transmitted by the *An. gambiae* complex. Mosquitoes were collected from 11 preliminary sites (Fig. 1B): seven urban (characterized by high population density, paved roads, markets) and four rural (subsistence farming, scattered housing, unpaved tracks). Two collection methods were employed per site: a) Early morning indoor resting collections of adult females (06:00–08:00) from 10 houses per site; b) Larval dipping from all accessible potential breeding sites within a 500 m radius. Specimens were stored in silica gel for transport. Raw data for the whole dataset are available in Supp. Table 3.

### 2.6. Environmental Noise Assessment

Ambient noise levels were measured at the centroid of each collection site using a Type 2 sound level meter (VLIKE LCD Digital Sound Level Meter). Measurements were performed every 30 minutes from 8:30 am to 5 pm over two consecutive days as a proxy for anthropogenic noise levels at each site, under the assumption that noise levels were maintained at sunset, when mosquito mating mostly occurs. Ten 30-second measurements of A-weighted equivalent continuous sound level were recorded per period at ∼1.5 m height. The mean sound levels were calculated for each site to characterize the chronic acoustic environment.

### 2.7. Statistical Analysis

Phonotaxis data (approach to speaker: yes/no) were analyzed using a logistic regression (binomial error distribution, logit link) including genotype, frequency, and their interaction as fixed effects. Post hoc comparisons of genotype within each frequency were performed using estimated marginal means with Tukey adjustment for multiple comparisons.

Mating success in the laboratory was analyzed using generalized linear mixed models (GLMMs) fitted with the glmer function from the lme4 package in R. The binary response variable (inseminated: yes/no) was modeled using a binomial error distribution with a logit link function. Fixed effects included acoustic condition (Low-noise: 70 dB vs. high-noise: 93 dB), male genotype (wild-type Tiefora vs. A296S-Hom), female genotype (wild-type Tiefora vs. A296S-Hom), replicate and the interactions between acoustic condition and male genotype and between acoustic condition and female genotype. Date was included as a random intercept to account for variation among experimental days. Post hoc comparisons of male genotype within each acoustic treatment were performed using estimated marginal means (emmeans), with Tukey adjustment for multiple comparisons. Each experimental run consisted of a single cage containing 25 males and 25 females of defined genotypes exposed simultaneously to one acoustic condition.

Differences in *Rdl* mutation frequencies between urban and rural sites were analyzed using Fisher’s exact test, with odds ratios and 95% confidence intervals calculated to assess effect sizes. For stratified analyses examining the association between *Rdl* genotypes and urban versus rural location while controlling for species and sex, the Cochran–Mantel–Haenszel test was used to obtain common odds ratios and 95% confidence intervals across strata. This approach accounts for potential confounding by species or sex and provides a pooled estimate of the urban–rural association.

Ambient noise levels between urban and rural sites were compared using Welch’s two-sample t-test. To model the relationship between environmental noise and the presence of resistant alleles, logistic regression was performed with *Rdl* mutant presence (yes/no) as the binary response variable and mean site noise level (dBA) as a continuous predictor.

All analyses were performed in R version 4.3.1.

## Acknowledgements

We thank the communities and authorities in Bangui for facilitating field collections. We acknowledge technical support from the teams at Institut Pasteur de Bangui and UCL. We would like to thank Bruno Gomes for manuscript revision and assistance from Picinali lab and Oliver J Turvey on measuring sound pressure levels. This work was supported by UK Research and Innovation under the Future Leaders Fellowship scheme (Grant reference MR/S015493/1 and MR/Y011732/1), a UCL Engagement Fund (2022/23 GEF), and a pilot grant from the Bill & Melinda Gates Foundation (INV-072333) to MA. This work also received funding from State Research Agency ATRAE Programme grant (ATR2023-145654), Plan Generación de Conocimiento 2024 (PID2023-146360OA-I00) to MA and the European Commission STOP-MATING consortium (101183033). MA is part of the CSIC’s Global Health Plaform (PTI Salud Global).

