## Supplementary Table 1 for "An auditory advantage of *Rdl*-resistant mosquitoes may promote its persistence in urban environments"

**Supplementary table 1: Summary of field collection sampling methods**

| **Site_area** | **Species** | **N** | **Larvae collections** | | | **Adult early morning collections** | | |
| --- | --- | --- | --- | --- | --- | --- | --- | --- |
|  |  |  | **Total** | Males | Females | **Total** | Males | Females |
| **Urban** | *An. coluzzii* | 90 | 80 | 48 | 32 | 10 | 3 | 7 |
|  | *An. gambiae* | 61 | 30 | 16 | 14 | 31 | 2 | 29 |
|  | *Hybrid* | 8 | 0 | 0 | 0 | 8 | 0 | 8 |
|  | Total Urban | 159 | 110 | 64 | 46 | 49 | 5 | 44 |
| **Rural** | *An. coluzzii* | 104 | 80 | 34 | 46 | 24 | 0 | 24 |
|  | *An. gambiae* | 189 | 160 | 57 | 103 | 29 | 0 | 29 |
|  | *Hybrid* | 21 | 20 | 6 | 14 | 1 | 0 | 1 |
|  | Total Rural | 314 | 260 | 97 | 163 | 54 | 0 | 54 |
