## Supplementary Table 2 for "An auditory advantage of *Rdl*-resistant mosquitoes may promote its persistence in urban environments"

| Site | location | Noise (mean ± SD) |
| --- | --- | --- |
| EC | Rural | 48.5 ± 3.53 dB |
| GB | urban | 54.1 ± 3.92 dB |
| IFB | rural | 51.6 ± 5.86 dB |
| Kol | urban | 72.5 ± 5.25 dB |
| Li | rural | 55.7 ± 8.01 dB |
| MBV | urban | 66 ± 4.78 dB |
| MR | urban | 71 ± 5.14 dB |
| OU | urban | 46.0 ± 4.73 dB |
| PK10 | urban | 51.8 ± 3.76 dB |
| PK6 | rural | 49.5 ± 4.72 dB |

**Supplementary table 2: Mean noise level in each site**
